# Enhanced cellular and transdermal delivery of the modified chromatin using gH625 cell-penetrating peptide

**DOI:** 10.1101/2025.09.11.675732

**Authors:** Xinghan Zhang, Yuxin Liang, Francesco Zonta, Jeong Hyeon Park

## Abstract

The transdermal drug delivery system has been used as a noninvasive route for drug administration, but often requires specialized equipment or invasive treatment. To examine recombinant synthetic chromatin as a non-invasive transdermal delivery vehicle, a modified histone H2A fused with a cell-penetrating peptide (gH625) was examined for its utility. Both *in vitro* porcine skin-based test and *in vivo* mice studies showed significantly enhanced passive skin penetration of gH625-modified chromatin compared to control groups. The gH625-H2A chromatin was examined as a delivery vehicle of EGF by displaying it at the carboxy terminus of histone H2B to enhance stability and skin penetration. The experimental results showed that the biological activities of EGF-displayed chromatin are comparable to those of the isolated EGF peptide in the HaCa-T cell line. The enhanced skin penetration capabilities of the gH625-H2A-modified chromatin offer great potential for developing non-invasive transdermal delivery of therapeutic proteins or peptides.

## INTRODUCTION

DNA is a highly negatively charged polyanionic polymer, which does not readily bind to or penetrate cell membranes and skin without a specialized uptake mechanism. Chromatin can be considered as a condensed polyanionic DNA polymer coated with cationic N-terminal histone tails ^1^. Interestingly, four core histones carry a protein transduction domain that facilitates membrane penetration. Additionally, these core histones, either individually or together, can condense DNA to facilitate DNA delivery into mammalian cells ^2-7^. More importantly, reconstituted chromatin enables DNA delivery to cells through an endocytosis-independent pathway ^8^. Collectively, histones show great potential as a non-viral DNA delivery vehicle due to their ability to condense nucleic acids, form stable nanocomplexes, and target specific cell delivery via histone modifications ^9-11^. However, the applications of histones for DNA delivery in mammals through systemic administration pose many challenges, such as ensuring histone-DNA complex stability, possible immunogenicity, and potential gene expression issues related to intracellular instability and histone-mediated gene repression ^9^.

The skin, as the largest organ by surface area in humans, is an attractive route for transdermal drug delivery due to its noninvasive nature and convenience ^12^. The transdermal route also minimizes nonspecific systemic side effects, making it highly appealing for both biopharmaceutical medicines and skincare applications ^13-16^. However, the epidermis, particularly the stratum corneum, serves as a highly effective barrier, making it difficult to deliver large biomolecules through passive topical application. Hydrophilic biomolecules with molecular weights over 500 Da generally require invasive administration or delivery enhancers such as microneedles and electroporation ^17^. Despite various physical and biochemical approaches, achieving efficient transdermal delivery remains a significant challenge ^18^. In connection, various cell-penetrating peptides (CPPs) have been explored for intracellular or skin delivery of therapeutic molecules by forming covalent or non-covalent complexes ^19-23^. One such peptide, gH625, derived from the glycoprotein H of herpes simplex virus type I (HSV-1), facilitates membrane fusion due to its hydrophobic amino acid residues and amphipathic α-helix formation ^24-26^. Previous studies have shown that gH625 peptide exhibits strong membrane insertion and penetration abilities ^24, 27-29^. Here, we demonstrate that human histone H2A, modified with gH625 peptide, enhances the cell penetration capabilities of histone H2A and chromatin. The chromatin assembled with gH625-H2A exhibited skin penetration superior to that of DNA and wild-type chromatin, suggesting its potential as a transdermal delivery vehicle for nucleic acids, small peptides, and enzymes without the need for physical or chemical interventions.

## RESULTS

### N-terminal extension of H2A with gH625 peptide facilitates cell penetration of the histone

The crystal structure of the nucleosome core particle shows that two N-terminal tails of histone H2A adopt unstructured domains with symmetric projections from the nucleosomal dyad axis ^30^. The gH625 peptide was directly fused to the N-terminus of histone H2A (Figure 1a), in which the gH625 region appears to adopt an α-helical conformation at the end of the flexible N-terminal tail of H2A (Fig. 1b). Previously, H2A was shown to possess the highest translocation ability across cell membrane among four core histones ^3, 5^. To compare the cell penetration ability of gH625-H2A with wild-type H2A in HeLa cells, the same amount of each biotinylated histone was directly added to the cell culture medium and the relative penetration efficiencies were visualized by Alexa Fluor™ 488 (Fig. 1c). While H2A wild-type treatment resulted in the modest visual fluorescence intensity, gH625-H2A treatment dramatically increased the fluorescence signal in cells, suggesting that more gH625-H2A histones penetrated the cells through the membrane (Fig. 1c, the images of Alexa Fluor™ 488, compare gH-H2A to WT H2A). The microplate readings of fluorescence signals indicated that gH625-H2A exhibited approximately six-fold higher fluorescence signals than wild-type H2A (Figure 1d). The result suggests that the fusion of the gH625 peptide to the N-terminus of H2A significantly enhances the intrinsic cell membrane penetration ability of H2A, achieving complete translocation within one hour (Supplementary Fig. 1a and 1b).

**Figure 1.**
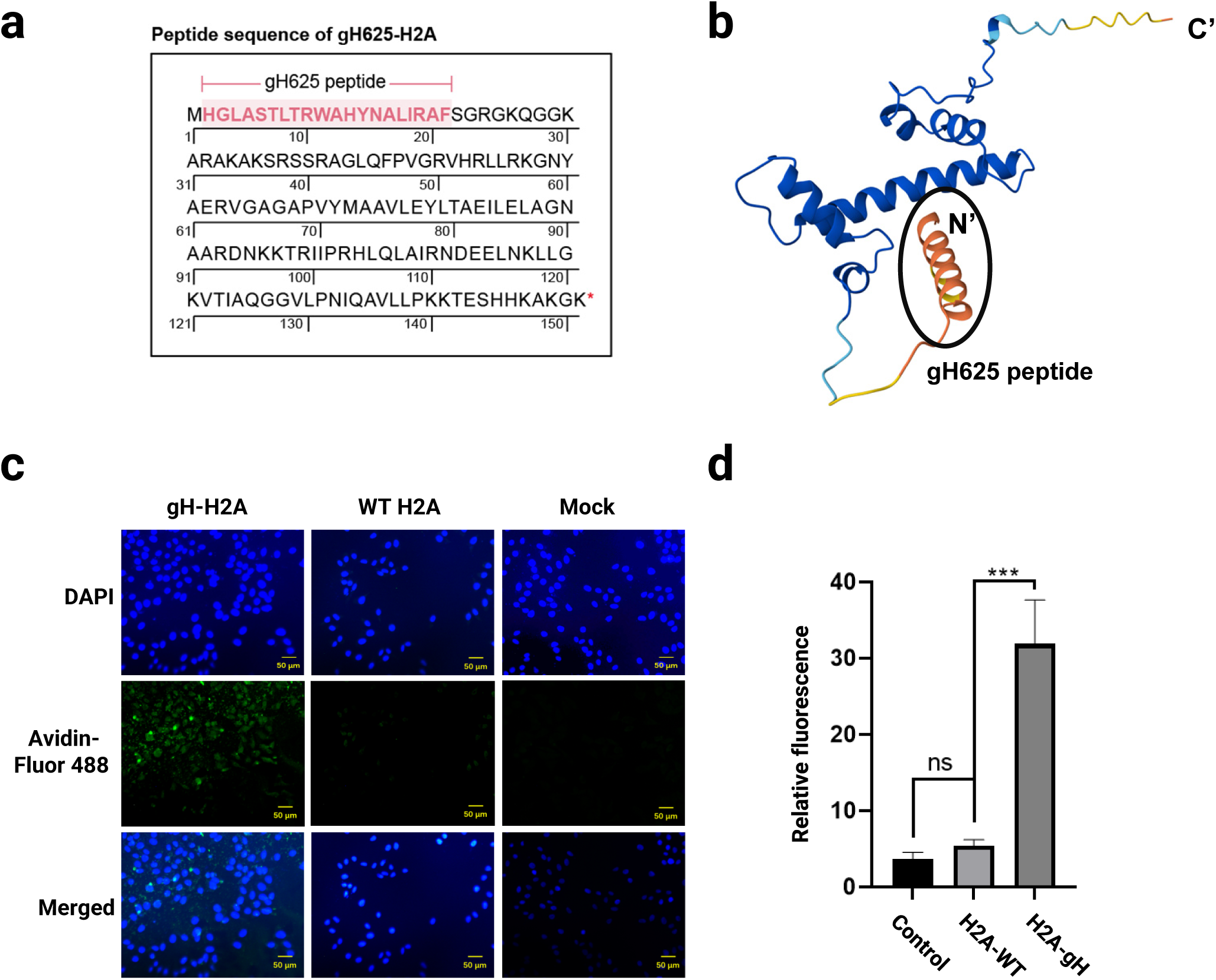
Structue of gH625-H2A and its cell membrane penetration. (a) Amino acid sequence of the gH625-H2A fusion protein, with the gH625 peptide highlighted in pink color. (b) AlphaFold 3 prediction of the gH625-H2A structure. (c) Cell membrane penetration assay of gH625-H2A in HeLa S3 cells. Representative images at 40x magnification with 1 s exposure are shown. (d) The microplate readings of fluorescence signals in panel (c) were plotted *ns*, non-significant.

### The gH625 peptide-displayed chromatin enhances the translocation of DNA across the membrane

To examine whether the gH625 peptide on the chromatin can enhance DNA delivery into the cells, the reconstituted histone octamers were prepared for *in vitro* chromatin assembly (Supplementary Fig. 1c). The chromatin was assembled using a biotinylated pUC19 plasmid containing 16 repeats of the 601-Widom nucleosome positioning sequence. The quality of the assembled chromatins was examined with MNase assay (Fig. 2a). While naked DNA mainly produced low molecular weight DNA fragments below 100 bp, chromatin assembled using histone octamers of either wild-type or gH625-H2A generated 200 bp-DNA ladders, suggesting that the chromatin mainly consists of physiologically space nucleosomes (Fig. 2a, compare lane 2 to lanes 4 and 6). These results indicate that the gH625 peptide extension at the N-terminus of H2A, as well as the biotin modifications of the plasmid DNA backbone, do not affect the quality of chromatin assembly.

**Figure 2.**
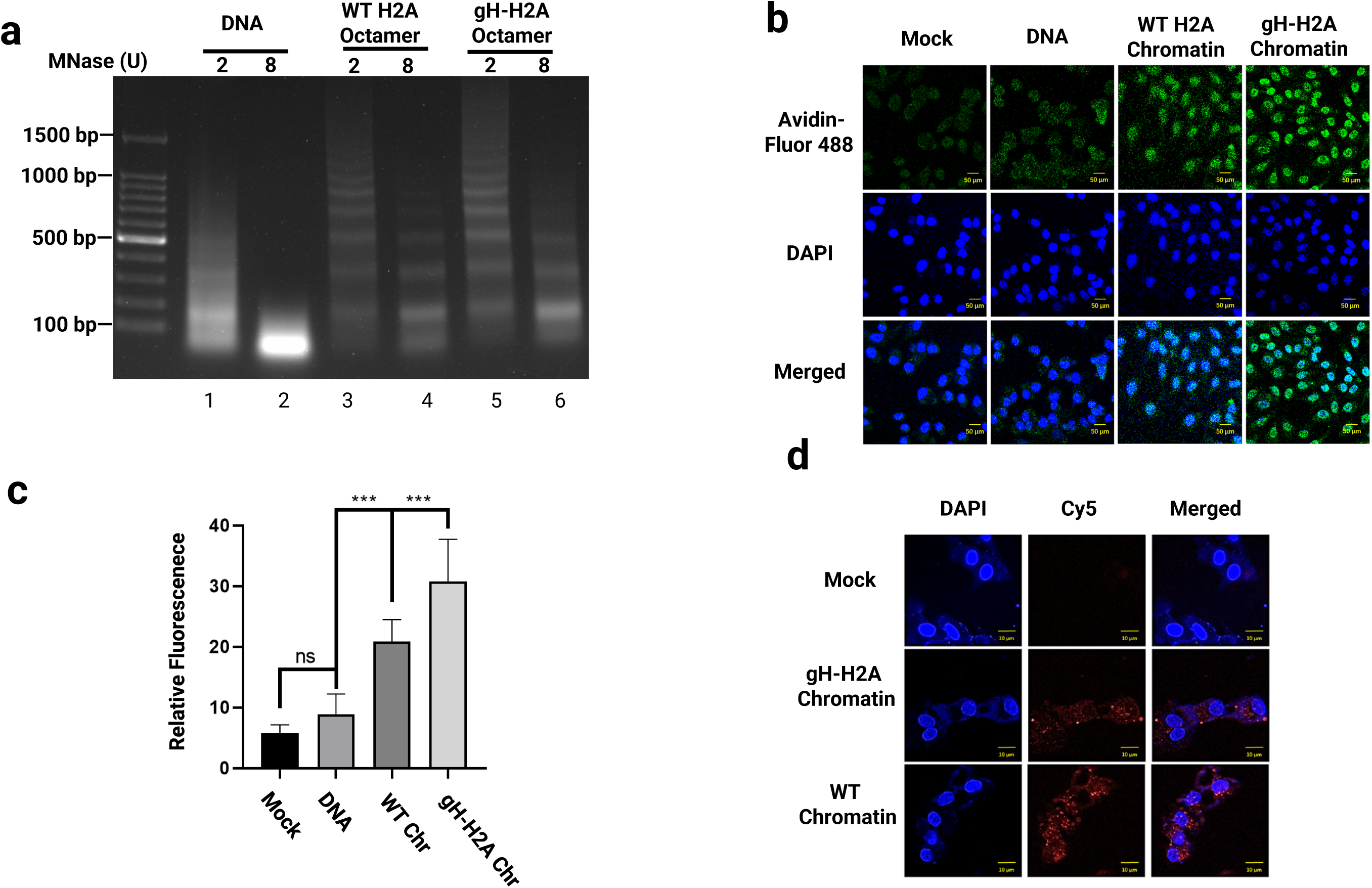
Cell membrane penetration assay of chromatin containing gH625-H2A. (a) MNase assay of the assembled chromatin containing gH625-H2A. (b) Cell membrane penetration assay of chromatin. Representative images at 40x magnification with 1 s exposure are shown. (c) The fluorescent intensities from panel (c) were analyzed by ImageJ software and the average fluorescent intensities were calculated and plotted. ns. non-significant. (d) Confocal image analysis of intracellular Cy5-labeled pUC19/16x601.

Similar cell penetration assays to those in Figure 1c were carried out using wild-type and gH625 peptide-displayed chromatins, respectively (Fig. 2b). The results show that gH625 peptide-displayed chromatin penetrated cells with higher efficiency than the wild-type chromatin (Fig. 2b, compare WT H2A chromatin to gH625-H2A chromatin in Alexa Fluor™ 488). Quantification of the fluorescence intensity in Figure 2b by Image J software confirms that wild-type chromatin exhibits intrinsic membrane penetration ability, and additional gH625 peptide on chromatin significantly increases plasmid DNA delivery efficiency across the cell membrane (Fig. 2c). To visualize the translocated plasmid DNA directly, DNA was prelabeled with Cy5 before chromatin assembly. HeLa cells grown on coverslips were treated with Cy5-labeled wild-type and gH-H2A chromatin, respectively and confocal fluorescence microscopic images were analyzed (Fig. 2d). Almost all cells treated with wild-type chromatin or gH-H2A chromatin displayed intracellular Cy5 signals. These results demonstrate that both wild-type chromatin and gH-H2A chromatin possess intrinsic membrane-penetrating capacity when directly incubated with cells, enabling them to enter cells by crossing the cell membranes.

### gH625 peptide-displayed chromatin improves skin penetration of DNA

Given the direct translocation mechanism of histones and chromatin across membranes, naked DNA and chromatin were examined for the direct skin penetration ability using a suckling pig skin-Franz cell system. Cy5-labeled plasmid DNA and its derived chromatin were applied three times on the pig skin at 24 h intervals in triplicate, and the frozen thin sections were analyzed under a fluorescence microscope (Fig. 3). The significantly higher fluorescence was observed in the stratum corneum after 24 h of exposure to gH625-H2A chromatin (Fig. 3a, 24 h column), which showed deeper penetration into the epidermis and dermis after 48 h and 72 h exposure. Wild-type chromatin also shows a better penetration than naked DNA, but its efficiency is significantly lower than gH625-H2A chromatin (Fig. 3b). The results demonstrate that the gH625 peptide on chromatin enhances chromatin penetration across skin barriers.

**Figure 3.**
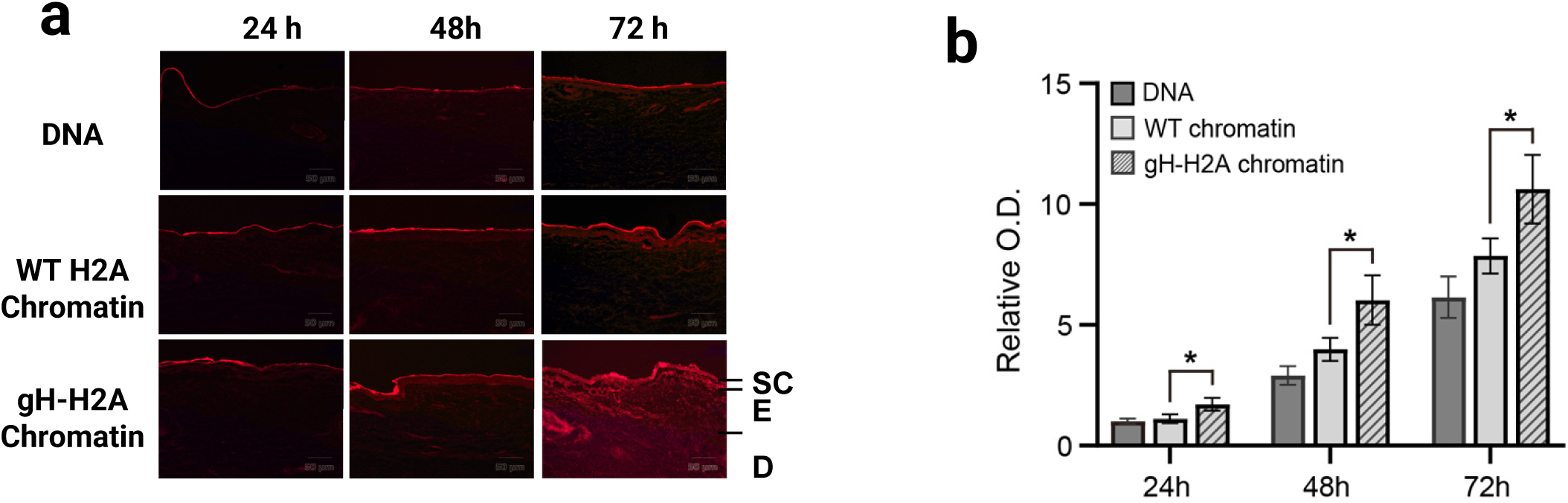
Skin penetration assay using a porcine skin-based Franz cell system. (a) Porcine skin penetration assay. Representative images of histological sections are shown. SC: stratum corneum; E: epidermis; D: dermis. (b) Fluorescence intensities at each time interval were measured in histological sections and the relative intensities were plotted. * p ≤ 0.05.

### Molecular dynamics (MD) simulations show improved stability of the EGF peptide when fused to the H2B protein in a nucleosome

The skin penetration test of gH625-H2A chromatin prompted us to examine whether gH625-H2A chromatin can be used as a delivery vehicle of large bioactive peptides that may reach the epidermis and stimulate cell growth and collagen secretion in keratinocytes. Since the H2B C-terminal tail extends from the histone core and remains relatively unstructured ^30^, human EGF peptide was fused at the C-terminus of human H2B histone using a short stiff linker. To assess protein stability of the EGF peptide alone or fused to chromatin, MD simulations were performed using a molecular model of a nucleosome containing a pair of H2B-EGF proteins. The initial model was generated using the AlphaFold 3 WebServer. As the linker between H2B and EGF contains only six amino acids, the EGF peptide cannot move far from the nucleosome (Fig. 4a and 4b).

**Figure 4.**
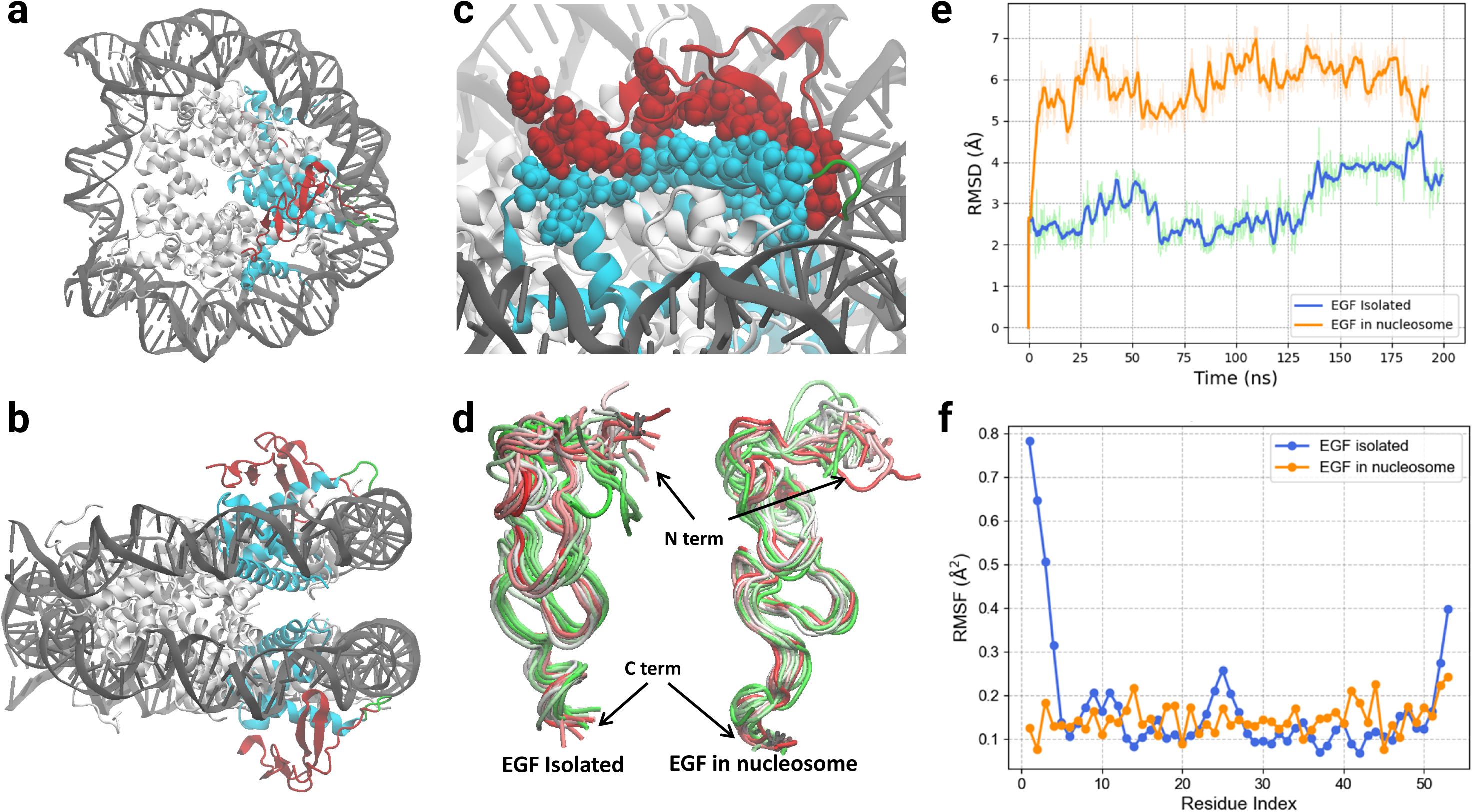
Molecular model of the H2B-EGF modified nucleosome. (a) and (b) Top and side view of the nucleosome respectively. The modified protein H2B part of the fusion protein is colored in cyan, while the EGF part is colored in red. The linker between the two is shown in green. The other proteins are white, while the DNA is gray. (c) The interaction network that stabilizes the position of the EGF portion of the fusion protein with respect to the rest of the nucleosome. (d) Configuration ensembles of the EGF peptide alone (left) or fused to the H2B peptide in the nucleosome (right) during a 200 ns MD simulation. Different configurations are color coded from red (beginning of the simulations) to green (end of the simulation). (e) and (f) Panel (e) and (f) compares respectively the RMSD and the RMSF of the EFG alone (blue line) or fused to the H2B protein (orange line).

Based on this model, it is evident that the EGF domain of the fusion protein is in close contact with the histone-fold domain of H2B. MD simulations suggest that significant conformational transitions of the EGF domain are unlikely to happen in physiological conditions, as the position of the EGF peptide linked to the H2B is rapidly stabilized (after 30 ns) through a network of interactions. These multiple interactions include the alpha helices of the H2B portion of the protein (residues 16-17, 60 to 69 and 77 to 88) and several sections of the EGF part (residues 101 to 103, 123 to 127, 140 to 144, 150) (Fig. 4c). In particular, the C-terminal region of EGF subdomain extends in a relatively short time and become stabilized by interaction with the H2B subdomain (Fig. 4d). By contrast, another simulation of a single EGF peptide in an aqueous environment shows a larger variety of possible configurations in the simulated timeframe (200 ns), especially on the N-terminus.

Analysis of the RMSD plot confirms these observations (Fig. 4e). The RMSD of the EGF subdomain fused to the H2B protein equilibrates after 30 ns, after a rapid increase due to the stretching at the level of the C-terminal domain. On the other hand, the EGF alone does not reach equilibration within the 200 ns observed timeframe, due to the large changes at the level of the N-terminal region. Consistent with this, calculation of the RMSFs over the trajectories after 30 ns clearly indicates the instability of the N-terminal domain and the whole peptide when the EGF is not linked to the H2B domain (Fig. 4f).

### EGF-displayed on gH625-chromatin shows a comparable biological activity of EGF in keratinocytes

To test biological activity of EGF fused on gH625-chromatin, the histone octamers containing modified histones of gH625-H2A and H2B-EGF were purified from *E. coli* expression system (Fig. 5a). The modified H2B-EGF carries an additional 6.8 kDa of EGF at the C-terminus and, once being incorporated into nucleosomes, enables two EGF peptides to be displayed per nucleosome on the chromatin structure. Chromatin assemblies *in vitro* using the octamers of modified histones were examined by the MNase assay (Fig. 5b). The chromatin assembly was successfully obtained with gH625-H2A/H2B-EGF octamers, although the quality of assembly based on the multiple DNA ladders appears to be deteriorated (Fig. 5b, lanes 1 and 2 versus lanes 3 and 4).

**Figure 5.**
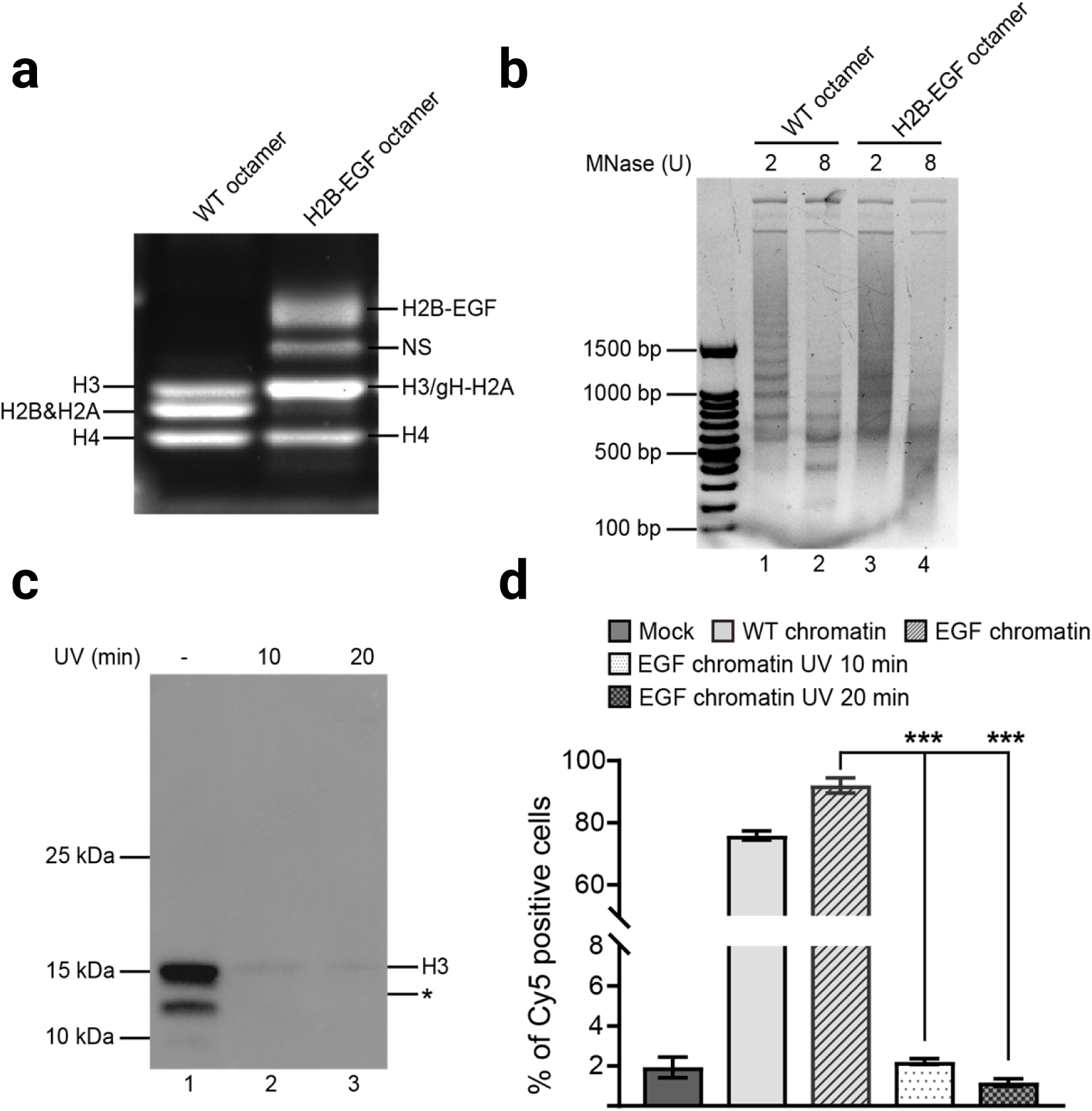
Assembly of EGF-displayed chromatin and UV-mediated crosslinking. (a) Octamers containing gH625-H2A and H2B-EGF were prepared and confirmed by SDS-PAGE followed by Coomassie Brilliant Blue G250 staining. NS, non-specific band. (b) MNase assay of the assembled chromatins. (c) Anti-histone H3 immunoblot analysis of UV crosslinked chromatins. *: non-specific band. (d) Cytometric analysis of Cy5 positive cells. The WT and EGF-displayed chromatins were directly applied to adherent HeLa cells and incubated for 3 h before cytometric analysis.

To assess the cell membrane penetration of EGF-displayed chromatin and the effects of UV irradiation-mediated crosslinking, the chromatin assembled from H2B-EGF histones was UV irradiated with different time periods (0, 10, 20 min) and analyzed by anti-H3 immunoblotting (Fig. 5c). The result shows a clear crosslinking effect with UV treatment longer than 10 min in which the free histone H3 released by SDS sample buffer was barely detectable in the UV-irradiated chromain (Fig. 5c, compare lanes 2 and 3 versus 1). The Cy5-labeled WT and EGF chromatins were then applied to HeLa cells and the Cy5-positive cell populations were quantified via flow cytometry. The EGF-displayed chromatins without crosslinking showed a better penetration efficiency than the WT chromatins. However, the crosslinked chromatin by UV irradiation appears to completely lose the penetrating ability of the cells, suggesting that the cell penetration activity might require a flexible and dynamic structure of the chromatin (Fig. 5d).

Although cell-penetrating chromatin is capable of crossing the cell membrane, we have previously shown that it can still effectively bind to cell surface membrane receptors ^11^. The biological activity of EGF-displayed chromatin was examined in terms of cell growth, collagen expression, and wound healing effects. The cell growth assay of EGF peptide and EGF-displayed chromatin showed that the treatment of EGF-displayed chromatin stimulated the cell growth of immortalized human keratinocytes, up to 150% compared to the mock control in the minimal growth medium (Fig. 6a). Assuming a maximum of 5% of the chromatin mass is from the EGF, EGF-displayed chromatin at 250 ng per ml is comparable to the known EGF peptide working concentration of 12.5 ng per ml in tissue-cultured cells (Supplementary Table 1). Therefore, the effective dose of EGF-displayed chromatin is comparable to the EGF peptide.

**Figure 6.**
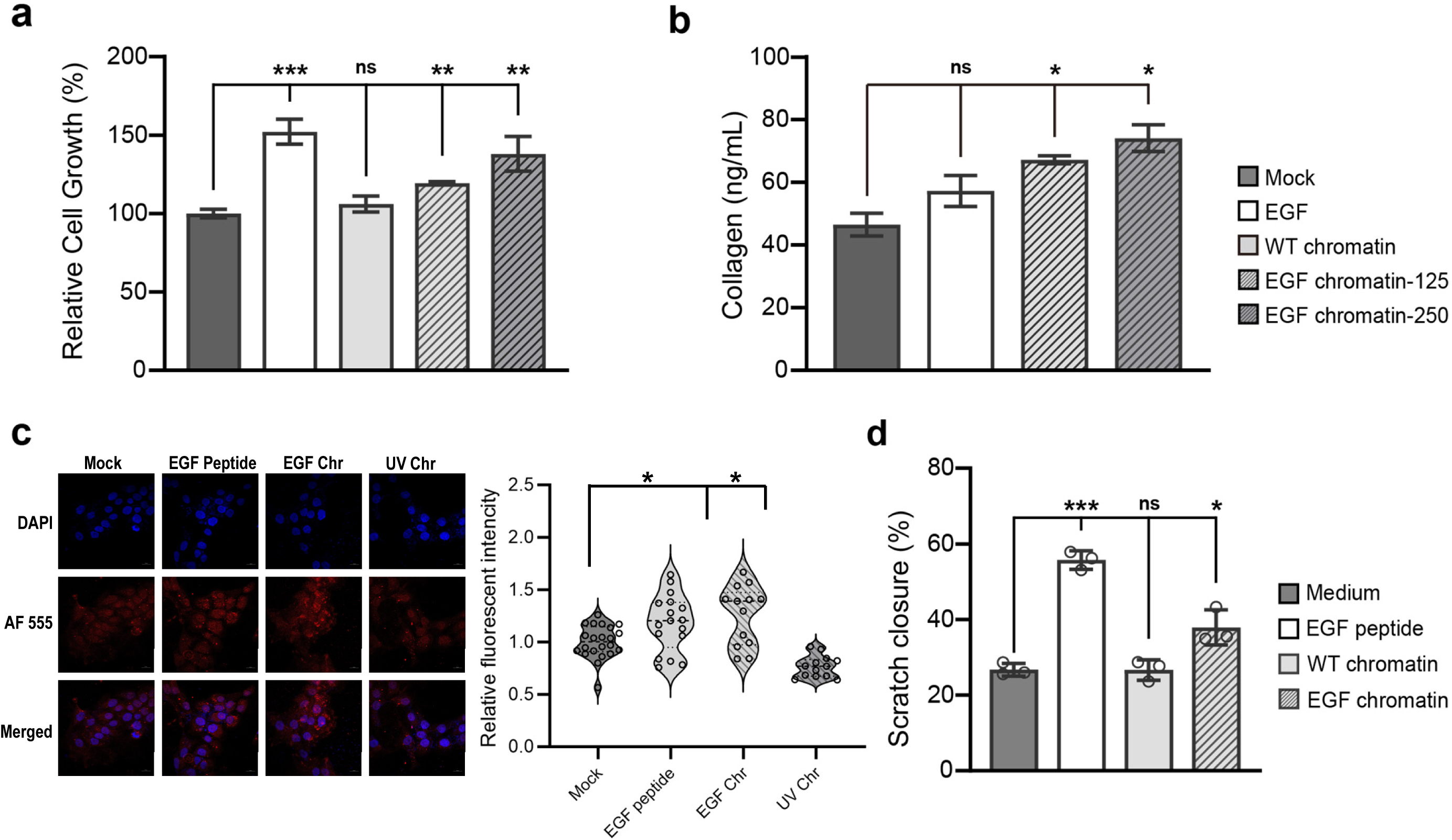
Cellular effects of EGF-displayed chromatin (a) Effects of EGF in HaCa-T cell growth. HaCa-T cells were treated with EGF (12.5 ng/ml), wild-type chromatin (250 ng/ml), and EGF-displayed chromatin (125 and 250 ng/ml) for three days. (b) Effects of EGF on the secretion of Collagen Type Iα1. The amount of collagen type Iα1 in the culture medium was quantified in triplicate using ELISA. (c) Effect of EGF-chromatin in stimulating collagene expression. HaCa-T cells were treated with EGF peptide (50 ng/mL), EGF-chromatin or UV-linked EGF-chromatin (500 ng/mL) in DMEM supplemented with 2% FBS for 48h. Representative confocal images at 40x magnification with 2s exposure are shown. Fluorescent intensities of each cell was analyzed by ImageJ software and plotted. * p ≤ 0.05. (d) Cell scratch assay. Quantification of scratch closure (%) in the cell scratch assay at t = 24 h is shown. Data are presented as the mean ± standard deviation (SD) from three independent experiments (n = 3).

EGF is known to slightly enhance collagen synthesis in type I collagen by acute exposure ^31^. The collagen secretion assay showed that EGF peptide appears to increase collagen type I secretion but its increase is not statistically significant to mock control (Fig. 6b). However, the treatment of EGF-displayed chromatin to HaCa-T cells induces a significantly higher concentration of collagen type I secretion into the medium (Fig. 6b). To further confirm the effect of EGF-displayed chromatin on collagen expression in keratinocytes, HaCa-T cells were stimulated with EGF-displayed chromatin, and intracellular collagen protein levels were assessed by immunostaining with an anti-collagen antibody (Fig. 6c). While basal collagen levels were not altered by UV-crosslinked chromatin, treatment of EGF peptide or EGF-displayed chromatin produces higher intracellular collagen immunofluorescence signals. To further investigate the effects of EGF-displayed chromatin on the migration capacity of HaCa-T cells, a scratch assay was performed with HaCa-T cells to evaluate wound healing. The closure of the scratch gap was monitored at 0 h and 24 h post-scratch. As shown in Fig. 6d, the scratch closure was significantly enhanced in cells treated with EGF-displayed chromatin compared to those treated with WT chromatin or medium alone. Taken together, the results in Figure 6 show that the significant modification of EGF as a chromatin fusion does not prevent the EGF from engaging cell-surface EGFR, suggesting that the structural integrity of the EGF domain and the biological activity appear to be maintained.

### EGF-displayed chromatin exhibits enhanced transdermal penetration in mouse skin

To evaluate the transdermal penetration ability of EGF-displayed chromatin *in vivo*, Cy5-labeled chromatinized DNA was applied to mouse skin, and the penetration depth was analyzed by H&E staining and fluorescence microscopy (Fig. 7a and 7b). While plasmid DNA exhibited shallow penetration, mainly reaching the epidermis and the epidermis–dermis junction at 24 hours, EGF-displayed chromatin emitted prominent red fluorescence and reached deep into the dermis with a granular distribution pattern. This distribution and signal intensity remained stable for EGF-displayed chromatin up to 72 hours, whereas a modest decrease was observed for plasmid DNA. The Cy5 fluorescence data and Image Pro Plus software were used to determine the maximum skin penetration depth (Fig. 7c). The gH625 chromatin group at 24 hours demonstrated the most significant average penetration depth (243.0 µm), which was significantly greater than that of DNA (181 µm).

**Figure 7.**
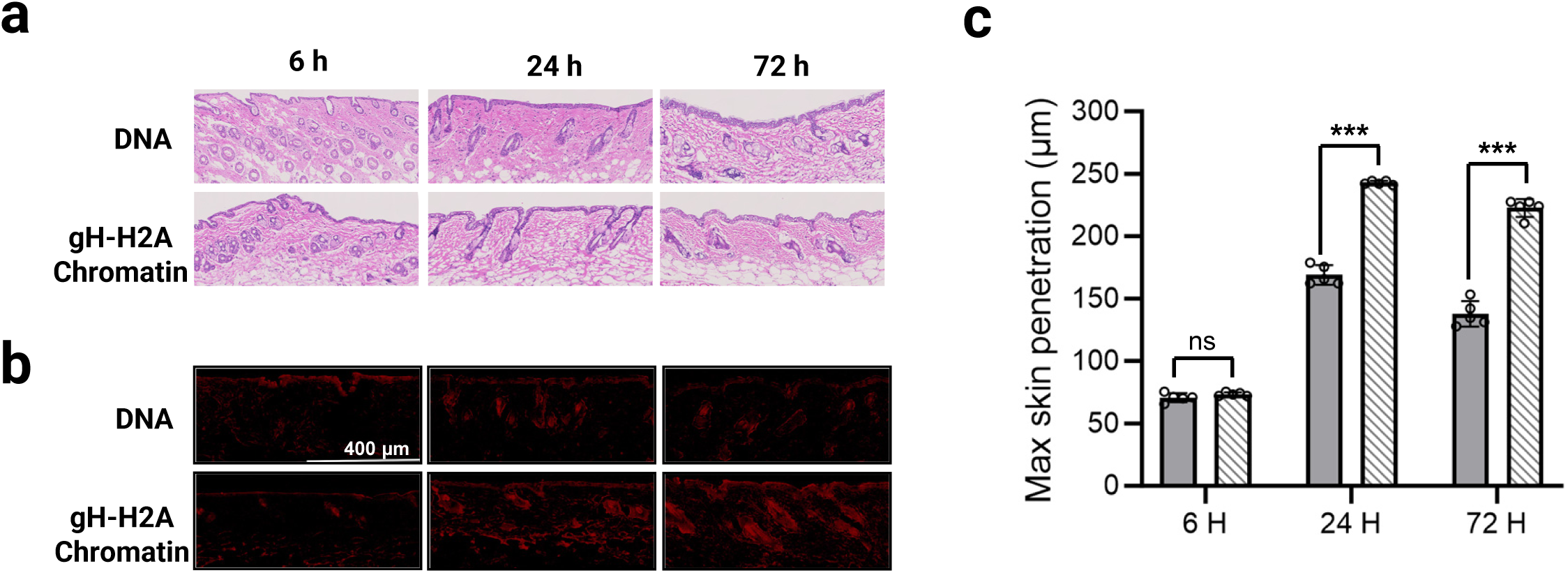
Transdermal penetration of EGF-displayed chromatin through mouse dorsal skin. (a) Hematoxylin and eosin staining of mouse dorsal skin for the skin structures, including the stratum corneum/epidermis and dermis. (b) Representative Cy5 fluorescence images of thin sections of mouse skin after DNA and chromatin applications. (c) Comparison of the skin penetration depth. Multiple fluorescence images were analyzed using Image-Pro Plus software to calculate the average penetration depth for each sample group.

Figure 7 clearly demonstrates the superior transdermal penetration ability of EGF-displayed chromatin compared to the control at 24 hours, consistent with the results of the porcine skin-based *in vitro* penetration study in Figure 3. Moreover, EGF-displayed chromatin did not permeate beyond the dermis into the subcutaneous area, indicating good biosafety for local transdermal drug delivery.

## DISCUSSION

The cell penetration ability of histones and chromatin has been previously reported but without elucidating a clear penetration mechanism ^9^. Given the fact that the efficiency of penetration through membrane is not affected by low temperature or common endocytic pathway inhibitors, it seems not to require a common endocytic entry pathway. A plausible model suggests that the hydrophilic nature, especially positively charged histone tails of chromatin, binds to negatively charged polar heads of phospholipid in the membrane, resulting in transient inverted micelles that would allow trapped molecules to cross the membrane ^32^. Consistent with the inverted micelle model of many fusogenic CPPs, hydrophobic residues of gH625 may cross the membrane by facilitating a transient pore formation and destabilization of the membrane bilayer ^26^. The application of gH625 to drug delivery has been extensively explored for many compounds ^26^. The ability of gH625, through direct membrane interaction, would contribute to increasing the intrinsic transduction ability of histones and chromatin across the membrane.

It is hard to imagine how hydrophilic chromatin with a molecular mass of over 10 MDa can pass the skin barrier passively. Three primary transdermal routes have been proposed for the peptide-based delivery: interstitial lipid barrier, trans-follicular path, and transcellular diffusion ^33^. The staining pattern of the mouse skin by gH625-displayed chromatin suggests that the molecules are initially absorbed through the hair follicle and gland area, but diffuse faster through transcellular diffusion. Future studies aiming to elucidate the precise penetration mechanism of chromatin through the skin remains to be explored.

EGF is one of the prototype growth factors that have been extensively used for the topical application of skin cosmetics and wound healing, based on its stimulatory effects on mitogenic regeneration activity and extracellular matrix formation. As for cosmetic EGF application, two major challenges are low transdermal delivery efficiency and protein instability ^34^. As proof of concept, our studies show that multiple arrays of EGF on chromatin could overcome these weaknesses. The 3-dimensional nucleosomal structure suggests that the H2B C terminus is exposed on the surface of the nucleosome core particle and commonly used for bigger peptide addition, such as GFP ^30^. Our studies also show that EGF-displayed chromatin induced a consistent increase in collagen expression, in which the biological activity of EGF appears to be stable for several months when stored at 4 °C. Given the observation of slower diffusion of chromatin through the skin and its molecular mass over 10 MDa, EGF-displayed chromatin is unlikely to induce systemic effects through the main bloodstream. Overall, chromatin as a transdermal delivery platform for cosmeceutical ingredient delivery is a promising platform in terms of safety, efficacy, and versatility by combining existing cosmeceutical peptides.

In conclusion, we show a potential exploitation of chromatin as a transdermal delivery vehicle by incorporating gH625-H2A histone. It could deliver a large peptide such as EGF or gene-encoding plasmid through the skin. Chromatin structure would stabilize delivered peptides and DNA from various proteases and DNase in the epidermis, and if degraded, facilitate the release of the delivered pharmaceutical and cosmeceutical peptides in the interstitial fluid and extracellular matrix area. Given minimal toxicity, immunogenicity, and greater structural flexibility and stability ^35^, further research on synthetic chromatin as a transdermal delivery vehicle will be useful for pharmaceutical and cosmetical peptide deliveries in the area of skin health and diseases.

## METHODS

### Purification of histone octamers and salt-gradient chromatin assembly

The pET3 expression vectors for histone genes have been extensively used for the expression and purification of wild-type histone ^36^. The addition of the gH625 and EGF to the histone genes was performed using DNA synthesis (GenScript Biotech Corp) from the sequences provided in the appendix. The preparation of histone octamers containing wild-type histones or a mixture of wild-type and modified histones was conducted as described previously ^35^. Large-scale chromatin assembly was performed using step-gradient dialysis, and the quality of the synthetic nucleosome arrays was assessed by a Micrococcal Nuclease (MNase) assay, as described previously ^35^.

### Cell penetration assay for histones and chromatin

Partially purified individual histones, wild-type H2A and gH625-H2A were conjugated with biotin molecules using a commercial kit, EZ-Link Pentylamine-Biotin (Thermo Scientific). For cell penetration assay, HeLa cells were seeded at a density of 10,000 cells per well in 96-well plates and incubated for 24 hours in growth medium. Either 5 μg of biotinylated wild-type H2A or gH625-H2A was applied directly to the growth medium and incubated for 1 h at 37 °C. The cells were thoroughly washed with PBS containing 0.5 mM MgCl_2_, fixed with 2% paraformaldehyde and permeabilized with 0.1% Triton X-100 in PBS for 15 min. After blocking with 10% bovine serum albumin (BSA) in PBS containing 0.5% Tween20, Avidin Alexa Fluor^TM^ 488 (Thermo Scientific, Product No. A21370) was applied into each well (0.5 μg per 100 μl of PBS) followed by DAPI staining. Alexa Fluor™ 488 alone, without histone addition, was used as a negative control. Quantification of histone penetration was performed by measuring fluorescent intensity with a plate reader in triplicate. For the cell penetration assay using biotin-labeled gH625-chromatin, HeLa cells were treated directly with chromatin in the same manner as the biotinylated histone treatment as described above. Fluorescence signals in cells were quantitatively and qualitatively visualized using a fluorescent microscope (Nikon inverted Microscope Ti-S), confocal fluorescent microscope (LSM 880, Zeiss), or a plate reader (Varioskan Lux, Thermo Scientific). Cell imaging data were analyzed using ImageJ. Statistical significance was indicated as follows: ns. non-significant; *: p < 0.05; **: p < 0.01; ***: p < 0.001.

### Investigation of Skin Penetration Using a Porcine Skin-Based Franz Cell System and live animal skin

Dorsal skin stored at -20 °C was thawed using deionized water and rinsed repeatedly with PBS buffer. The pig skin was fixed between the donor and receiving chambers of the Franz cell, ensuring the stratum corneum faced the donor chamber. A total of 7.0 ml of receiving solution was added to the receiving chamber, followed by 1.0 ml of PBS introduced through a sampler to ensure close contact with the dermis. Each 0.5 µg DNA-equivalent sample was applied to the skin surface in the donor chamber, covering an effective area of approximately 3.14 cm², and spread radially from the center. The chamber solution was stirred at 300 rpm in a water bath maintained at 32 °C to facilitate absorption. The skin samples were collected at designated time points (24 h, 48 h, and 72 h) and washed five times with PBS. The samples were fixed in 4% paraformaldehyde for at least 24 h before frozen section analysis.

Animal studies were performed in compliance with guidelines set by Guangdong University of Technology’s Institutional Animal Care and Use Committee (#TRCW0606). 8-week-old female KM mice were used for all the in vivo experiments. Each Cy5-labeled DNA and chromatin solution (10 µg/ml) was evenly applied to the shaved dosal skin area and the skin samples were harvested at 6 h, 24 h, and 72 h. After embedding and sectioning, H&E staining was used to distinguish skin layers, and inverted fluorescence microscopy was used to observe distribution and depth of penetration of Cy5-labeled DNA. The penetration depth was quantified with Image pro plus software.

### EGF activity assays in human keratinocytes

For cell proliferation assay, 10,000 HaCa-T cells were plated in a 96-well plate and treated with a range of reagents on the same day. These treatments included varying concentrations of the EGF peptide and EGF-displayed chromatin. Following a two-day incubation period, cell growth was evaluated using the Cell Counting Kit (CCK-8) (Beyotime, China) according to the product manual. To analyze the collagen secretion in HaCa-T cell cultures over a 48-hour period, the cells were treated with the EGF peptide and EGF-displayed chromatin in the same manner as in the cell growth assay. The culture media from the 96-well plate were collected in triplicate at 48 h. A human COL1α1 ELISA Kit (Elabscience, China) was used for collagen quantification according to the product manual. To assess intracellular collagen expression, HaCa-T cells were grown on the slide and stimulated by adding EGF (50 ng/ml) and chromatin (500 ng/ml) for two days. Washed cells were fixed by 4% paraformaldehyde and permeabilized by 0.2% Triton X-100 in PBS. Cells were incubated by anti-collagen I antibody (Abcam, ab34710, 1:200 dilution) in the blocking buffer (PBS + 5%BSA + 0.5% Tween-20) overnight at 4 C. Washed cells were stained with Texas red goat anti-rabbit secondary antibody for 1hr at room temperature. The stained samples were counterstained with DAPI soltion and visualized under a fluorescence microscope. For the wound healing assay (cell scratch assay), artificial wounds were introduced into the HaCa-T cell monolayers using a 200 μL pipette tip. The cells were then washed three times with PBS to remove detached cell debris. The wound margins were imaged at the initial time point (t = 0 h). The cells were then treated with basal medium (DMEM with 1% Fetal Bovine Serum), EGFP peptide (12.5 ng/ml), WT-chromatin (500 ng/ml), or EGF-chromatin (500 ng/ml) for a duration of up to 24 h. The same wound margin fields were imaged at subsequent time points (t = 12 h and t = 24 h) using a Ti-S fluorescence microscope (Nikon) equipped with NIS-Elements imaging software. The percentage of wound area covered by the cells was quantified using ImageJ software.

### Molecular modeling and Molecular Dynamics of the H2B-EGF modified nucleosome

Molecular Dynamics (MD) simulations on this H2B-EGF-nucleosome model and the EGF peptide alone, were performed using protocols as described previously ^37^. Each system was solvated with TIP3P water, containing Cl- and K+ ions at a concentration of ∼ 0.15 M to mimic physiological ionic conditions. The total atom count after solvation was approximately 4 × 10^5^ for the H2B-EGF-nucleosome, and 6 × 10^4^ for the EGF peptide alone. The molecular systems underwent energy minimization followed by 1ns pre-equilibration simulations in the NTV ensemble. Finally, a 200 ns production run was performed in the NPT ensemble, with a 2 fs timestep. Long-range electrostatics were handled using the particle mesh Ewald method ^38^, with a 12 Å cutoff for Lennard–Jones interactions and a switching function starting at 10 Å. Temperature was maintained at 310 K using the V-rescale thermostat with a 1 ps coupling constant, and pressure was controlled at 1 bar using the Parrinello-Rahman barostat with a 5 ps coupling constant ^39^. The LINCS algorithm was used to constrain hydrogen-containing bonds. All simulation were performed in Gromacs^40^, with Amber ff19SB Force Field ^41^. Figures and part of the analysis was done using VMD ^42^.

## Acknowledgments

We thank Guantong Chen, Sam Shuang, Wenjing Xu, and Jianan Luo for their technical assistance for protein purification and FACS analysis and Qiujie Zhu for technical support in MD simulations. We thank Guangdong Technical University’s animal facility for assessing animal skin penetration activity. This research was partially supported by XJTLU Postgraduate Research Scholarship to Xinghan Zhang and Yuxin Liang, and by XJTLU Research Development Fund (RDF-23-01-026) to FZ.

## Author Contributions

JHP conceived and designed the experiments. XZ and YL performed the experiments. JHP, XZ, and YL analyzed the data and wrote the paper. FZ generated the nucleosome model, performed MD simulations, analyzed the corresponding data and wrote the part. All authors discussed the experimental results and edited the paper.

---

**Supplementary Figure 1.**
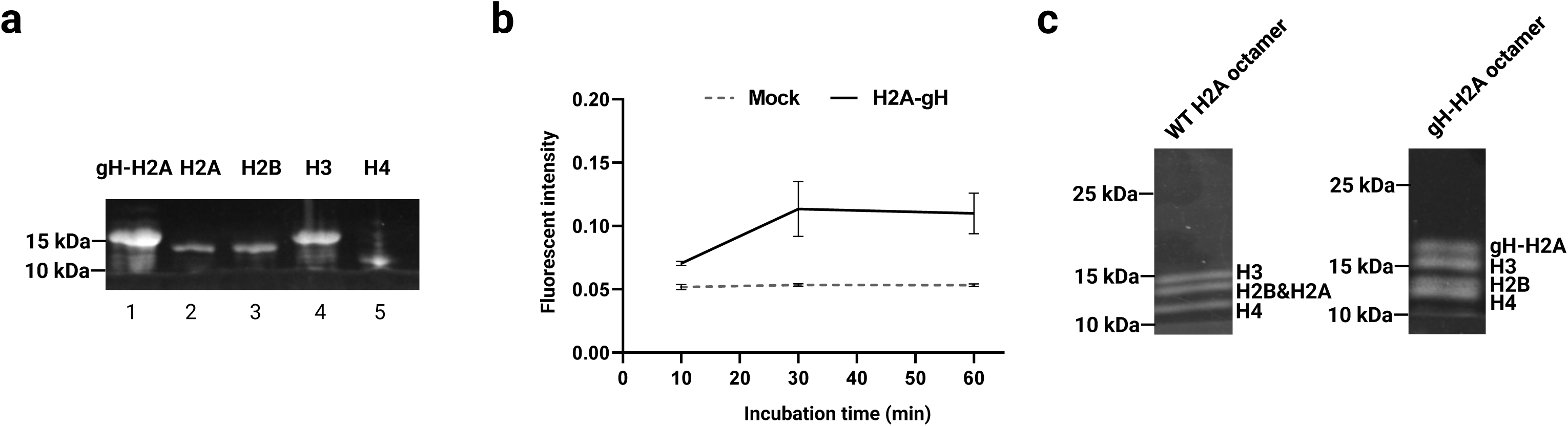
(a) Individual histone preparations from *E. coli* BL21. The purified histones were confirmed by SDS-PAGE and Coomassie Brilliant Blue G250 staining. (b) Time course of cell membrane penetration by gH625-H2A. (c) Histone octamers containing wild-type H2A or gH625-H2A were prepared and confirmed by SDS-PAGE and Coomassie Brilliant Blue G250 staining.

**Supplementary Table 1.** Biophysical properties of histone variants.

| **Protein name** | **Protein ID** | **Length** | **MW (KDa)** | **PI** | **Extinction coefficient** | **Instability Index** |
| --- | --- | --- | --- | --- | --- | --- |
| H2A | Human 2-A<br>(NP_003507.1) | 132 | 14.1 | 10.9 | 4470 | 53.96 |
| H2B | Human H2-E<br>(NP_003519.1) | 128 | 13.9 | 10.3 | 7450 | 46.08 |
| H3 | Human H3.2<br>(NP_066403.2) | 138 | 15.3 | 11.3 | 4470 | 47.55 |
| H4 | Human H4<br>(NP_003539.1) | 105 | 11.3 | 11.4 | 5960 | 43.5 |
| gH625-H2A | HSV_gH625-H2A | 150 | 16.3 | 11.1 | 11460 | 53.24 |
| H2B-EGF | H2B-EGF | 185 | 20.7 | 9.6 | 25900 | 48.26 |

Parameters are calculated using Expasy Protparam (Gasteiger E., Hoogland C., Gattiker A., Duvaud S., Wilkins M.R., Appel R.D., Bairoch A.; Protein Identification and Analysis Tools on the Expasy Server; (In) John M. Walker (ed): The Proteomics Protocols Handbook, Humana Press (2005). pp. 571-607)

